# Bacterial immune evasion proteins: The therapeutic potential of CHIPS

**DOI:** 10.1101/2020.02.25.964908

**Authors:** Angelino T. Tromp, Yuxi Zhao, Ilse Jongerius, Erik C.J.M Heezius, Pauline Abrial, Maartje Ruyken, Jos A. G. van Strijp, Carla.J.C. de Haas, András N. Spaan, Kok P.M. van Kessel, Thomas Henry, Pieter-Jan A. Haas

## Abstract

Bacterial pathogens have evolved to secrete strong anti-inflammatory proteins that target the immune system. It was long speculated whether these virulence factors could serve as therapeutics in diseases in which abnormal immune activation plays a role. We adopted the secreted Chemotaxis Inhibitory Protein of Staphylococcus aureus (CHIPS) as a model virulence factor-based therapeutic agent for diseases in which C5aR1 stimulation plays an important role. We show that administration of CHIPS in human C5aR1 knock-in mice successfully dampens C5a mediated neutrophil migration during immune complex-initiated inflammation. Subsequent CHIPS toxicology studies in animal models were promising. However, during a small phase-I trial, healthy human volunteers showed adverse effects directly after CHIPS administration. Subjects showed clinical signs of anaphylaxis with mild leukocytopenia and increased C-reactive protein concentrations, suggesting an inflammatory response, which are believed to be related to the presence of relatively high circulating anti-CHIPS antibodies. Even though our data in mice shows CHIPS as a potential anti-inflammatory agent, safety issues in human subjects temper the use of CHIPS in its current form as a therapeutic candidate. The use of staphylococcal proteins, or other bacterial proteins, as therapeutics or immune-modulators in humans is severely hampered by pre-existing circulating antibodies.

## Introduction

The human immune system is a well-balanced and effective network of cells, tissues and organs, and plays a crucial part in the continuous fight against invading microbes. On the other hand, the survival of microbial pathogens dependents on their ability to withstand attacks by the immune system. Successful pathogenic bacteria have coevolved with the host and acquired complex methods of subverting and suppressing the immune system. The deployment of strong and specific immune modulatory proteins by bacteria have shown to be effective immune suppressors *in vitro* and *in vivo* in mice. Considering that abnormal or excessive activation of the immune system can lead to inflammatory diseases, it was long speculated whether these bacterial virulence factors could serve as anti-inflammatory therapeutics in conditions in which undesirable immune activation plays a role. Over the years, studies have suggested the therapeutic potential of various bacterial proteins that normally play a role in immune evasion. However, as bacterial-derived proteins will induce antibody responses, it remains enigmatic whether these proteins can indeed serve as means for anti-inflammatory treatments in humans. Examples of known pathogenic bacteria that secrete immune-evasion proteins are *Streptococcus pneumoniae*, *Pseudomonas aeruginosa*, *Neisseria gonorrhoeae* and *Listeria monocytogenes*. However, secreting more than 35 immune-evasion molecules [1], *Staphylococcus aureus* is the text-book example of immune evasion by bacteria.

*Staphylococcus aureus*, a common colonizer of human skin an nose as well as a human pathogen, has evolved to secrete an arsenal of virulence factors that target the human immune system [2]. One extensively described and well-studied *S. aureus* virulence factor is the Chemotaxis Inhibitory Protein of *Staphylococcus aureus* (CHIPS). CHIPS binds to the N-terminus of human C5aR1 with high affinity (*K*_d_=1.1nM) and functionally blocks the interaction with C5a, thus preventing C5aR1 stimulation and antagonizing chemotaxis [3–5]. Besides playing a role in chemotaxis as a response to microbial invasion, C5aR1 is involved in a variety of other inflammatory processes. Upregulation of C5aR1 on internal organs during the onset of sepsis, together with the excessive release of C5a, was proposed to lead to multi organ failure and death in rats [6,7]. Blockade of C5aR1 with polyclonal anti-C5aR1 antibodies was protective and increased survival in animal sepsis model [6]. C5a and C5aR1 have also been described to be involved in disease processes such as ischemia-reperfusion injury, rheumatoid arthritis, asthma, immune complex diseases, neurodegeneration and Alzheimer’s disease [8–10]. Targeting of the C5aR1 has shown to be beneficial in some of these disease processes in animals, emphasizing the relevance of the C5aR1 as a therapeutic target [11–18].

The properties of CHIPS to inhibit the human C5aR1 with high specificity and affinity makes it an example of a promising anti-inflammatory drug candidate in diseases in which C5aR1 stimulation plays an important role. Previous studies have shown that the antagonistic activity of CHIPS on mouse C5aR1 is 30-fold lower compared to human C5aR1 expressing cells [3]. This human specificity of CHIPS has hampered the assessment of CHIPS *in vivo* during inflammation and infection. Here, we report the application of a transgenic human C5aR1 knock-in mouse (hC5aR1^KI^) to assess CHIPS as a model anti-inflammatory compound in C5aR1-mediated diseases. Furthermore, we investigate the safety and efficacy of CHIPS in a phase-I, randomized, double blind, placebo-controlled study in humans.

## Results

### CHIPS binds hC5aR1 ^KI^ murine neutrophils and inhibits stimulation by murine C5a

In order to validate the suitability of our hC5aR1^KI^ mouse [19] as a model to evaluate CHIPS *in vivo*, we first assessed the activity of CHIPS on hC5aR1^KI^ murine neutrophils. To this end, the binding of CHIPS on bone-marrow derived hC5aR1^KI^ murine neutrophils was determined and compared to human neutrophils isolated from peripheral blood. We confirmed that CHIPS binds to hC5aR1^KI^ murine neutrophils at comparable levels as human neutrophils (Figure 1a). To further assess the activity of CHIPS, inhibition of hC5aR1 was assessed on human and hC5aR1^KI^ murine neutrophils. Wild-type (*wt*) murine neutrophils respond normally to mC5a but CHIPS is ineffective in inhibiting mC5a-mediated Ca-mobilization on these mC5aR expressing cells (Figure 1b). Correspondingly, CHIPS inhibition of mC5a mediated Ca-mobilization of bone-marrow derived hC5aR1^KI^ neutrophils reflected that of human neutrophils isolated from peripheral blood (Figure 1b). Hereby, we confirm the binding and inhibition of hC5aR1^KI^ murine neutrophils by CHIPS, proving that our hC5aR1^KI^ mouse is a suitable model to assess CHIPS activity *in vivo*.

**Fig. 1:**
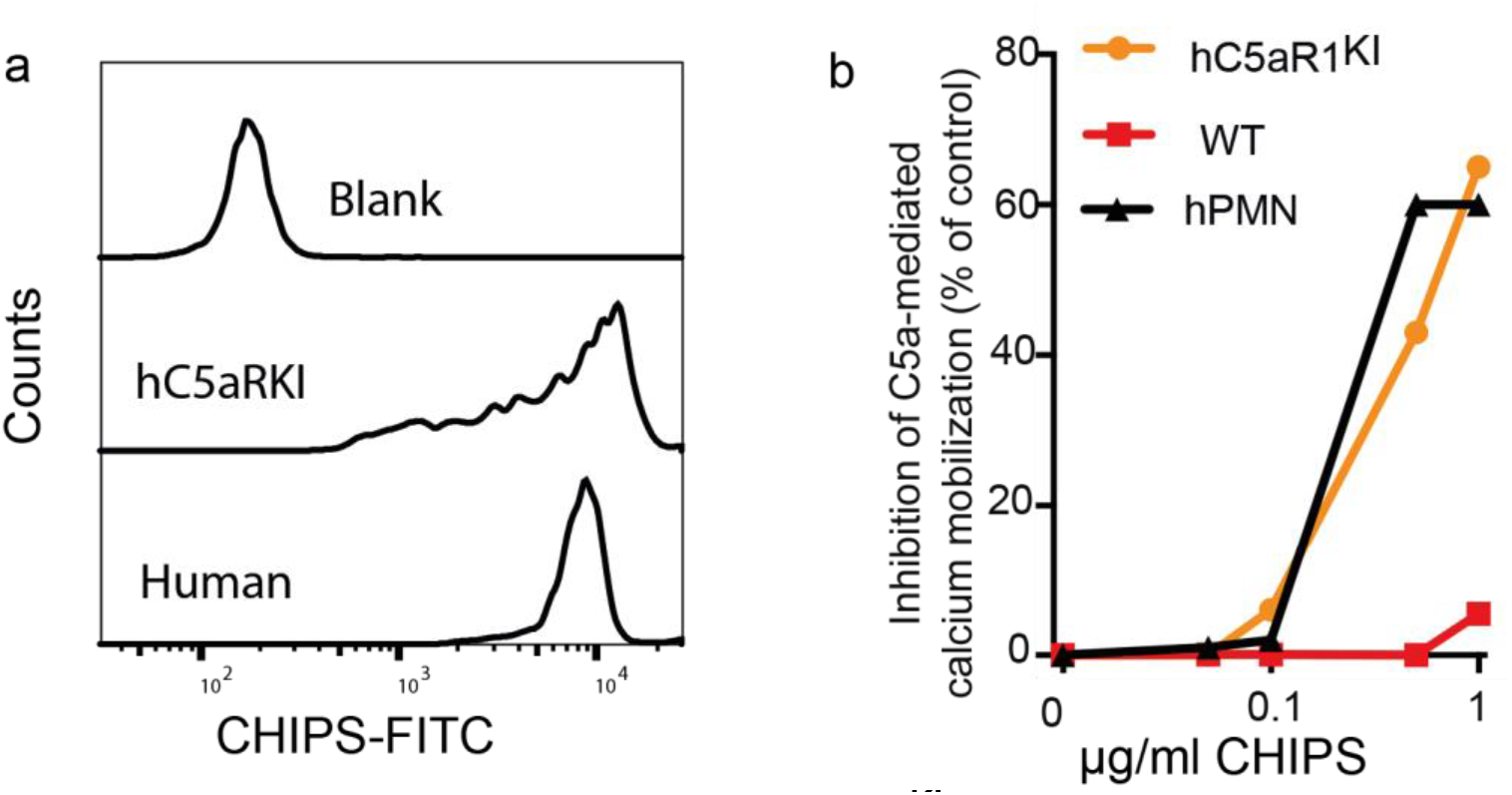
CHIPS binds and inhibits hC5aR^KI^ murine neutrophils comparable to human neutrophils. Quantification of hC5aR1 expression in hC5aR1^KI^ mice showed similar expression levels compared to human leukocytes[19]. Furthermore, hC5aR1^KI^ murine neutrophils responded normally to both murine C5a (mC5a) and human C5a as measured by calcium mobilization[19].**a,** hC5aR^KI^ bone marrow neutrophils and human blood neutrophils were isolated and incubated with 3μg/ml his-tagged CHIPS followed by anti-his-FITC antibodies. Cells were analysed by flow cytometry and the FITC fluorescent signal depicted as histograms. **b,** As our hC5aR1^KI^ murine model generates mC5a, the assessment of CHIPS inhibition was performed by mC5a stimulation. Bone marrow neutrophils of hC5aR^KI^, wild-type mice and human neutrophils were pre-incubated with CHIPS at the indicated concentration and subsequently stimulated with murine C5a (10-8M). The basal fluorescence level was first measured for each sample before the addition of murine C5a. The C5a-mediated calcium influx was analysed by flow cytometry using FLuo-4AM. The average FLuo-4AM fluorescent signal was used to calculate CHIPS mediated inhibition of C5a responses. One experiment representative of two independent experiments is shown.

### CHIPS inhibits C5aR mediated neutrophil migration in vivo

To assess the *in vivo* therapeutic potency of CHIPS, the immune complex-mediated Arthus reaction model [20,21] was used in hC5aR1^KI^ mice. The resulting inflammatory response and neutrophil recruitment in the Arthus reaction is mainly C5a mediated. By simultaneously administering ovalbumin intravenous (i.v.) and rabbit anti-ovalbumin IgG intraperitoneal (i.p.), an immune complex mediated type 3 hypersensitivity reaction is induced that leads to activation of the complement system and the generation of C5a [20,21]. An Arthus reaction was successfully induced in hC5aR1^KI^ mice as reflected by the influx of neutrophils to the peritoneal cavity (Figure 2a). Administration of CHIPS reduced the number of neutrophils recovered from the peritoneal cavity of hC5aR1^KI^ mice (Figure 2a). Some mice that received CHIPS showed suboptimal inhibition of neutrophil migration, whereas a single mouse showed no evident decrease in neutrophils recovered compared to untreated mice (Figure 2a).

**Figure 2:**
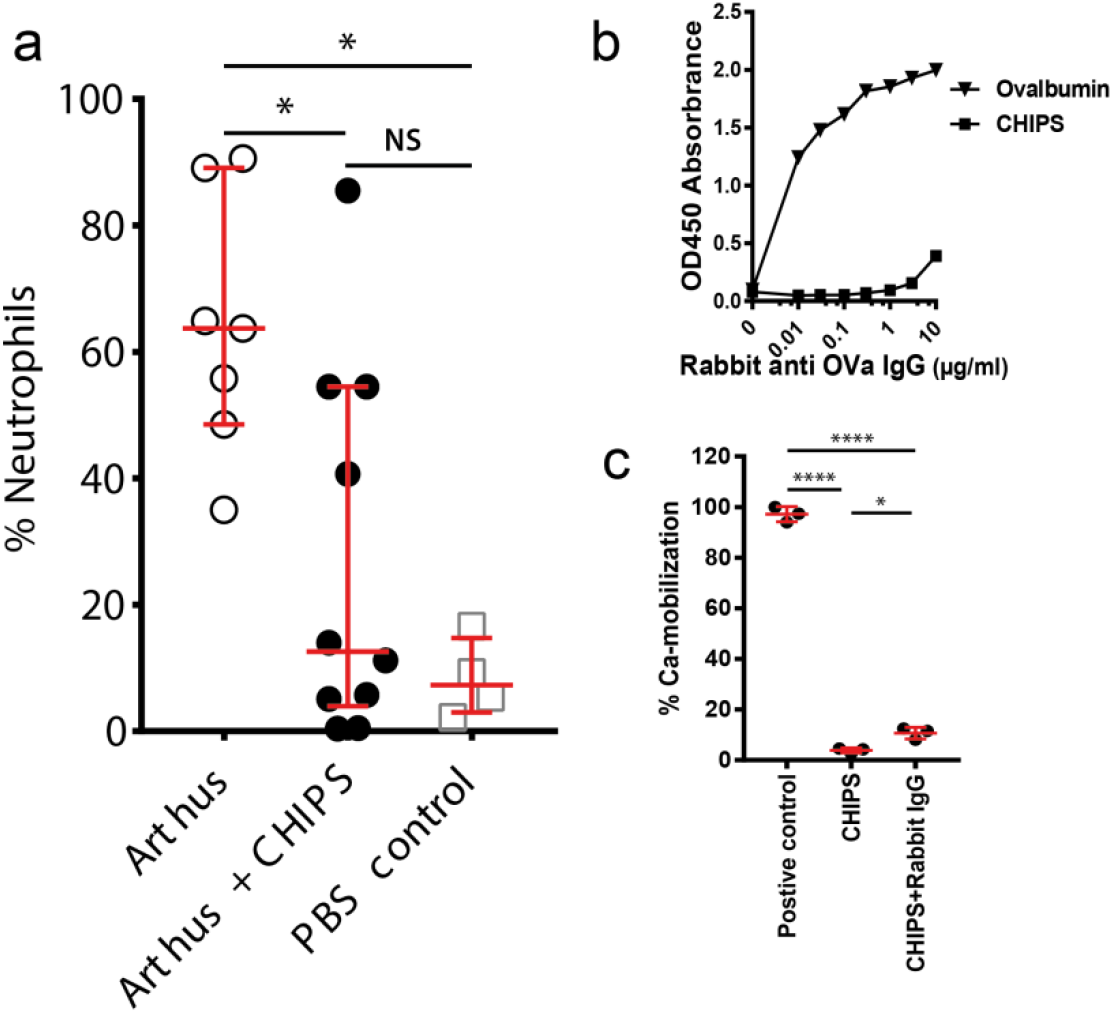
CHIPS inhibits neutrophil migration *in vivo*. **a,** 60μg CHIPS (*n*=10) was injected i.p., and together with ovalbumin i.v. in hC5aR^KI^ mice 30 minutes prior to inducing the Arthus reaction. Samples were compared to mice that did not receive CHIPS (*n*=7). Control mice (*n*=4) received PBS i.v. and i.p.. Peritoneal cavity lavage was performed 6-hours post Arthus induction. Percentage neutrophil influx was analysed by flow cytometry by gating on CD45^+^GR-1^+^F4/80^−^ population and depicted as percentage of total leukocytes (CD45^+^) retrieved after peritoneal lavage. All groups consisted evenly out of female and male mice. Combined data from 2 independent experiments shown. **b,** the presence of anti-OVA and anti-CHIPS antibodies in the rabbit anti-OVA IgG fraction was determined by ELISA. **C)** To detect neutralizing anti-CHIPS antibodies in the rabbit anti-OVA IgG, CHIPS (500ng/ml) was incubated with 10μg/ml rabbit anti-OVA IgG or PBS. Subsequently, Fluo-4AM labelled human PMNs were incubated with CHIPS/Rabbit IgG or CHIPS/PBS and challenged with human C5a. Ca-mobilization was determined via flow cytometry and normalized to human PMNs that did not receive CHIPS. Significance was calculated using ANOVA, and when needed, followed by Kruskal-Wallis post-test for multiple comparison and displayed as *\*P*<0.05,*\*\*\*\*P*<0.0001 and NS for not significant.

As *S. aureus* also colonizes rabbits [22], it is possible that the rabbit anti-ovalbumin IgG fraction used to induce formation of immune complexes also contains specific antibodies against CHIPS with potentially neutralizing capacities. To this end, we determined the presence of anti-CHIPS antibodies in the rabbit anti-ovalbumin IgG used. Although the rabbit IgG fraction did contain very low levels of anti-CHIPS antibodies (Figure 2b), the presence of these anti-CHIPS antibodies only slightly neutralized CHIPS *in vitro* and evidently did not neutralize CHIPS *in vivo* (Figure 2c, a). Taken together, our investigations demonstrate the therapeutic potential of CHIPS by inhibiting C5a-mediated neutrophil migration *in vivo* in hC5aR1^KI^ mice after inducing an Arthus reaction.

### CHIPS in human volunteers

*S. aureus* is commonly present as a commensal bacterium in humans and the *chp* gene is present in the majority of *S. aureus* strains. Consequently most, if not all humans, carry pre-existing anti-CHIPS antibodies [23–26]. These anti-CHIPS antibodies present in human sera have been shown to interfere with CHIPS function *in vitro* [23]. As a consequence, the presence of anti-CHIPS antibodies could neutralize CHIPS or induce an antibody-mediated immune reaction *in vivo,* hampering CHIPS function. To have an indication how subject titers relate to the general population, anti-CHIPS IgG titers were determined in sera collected from 168 human volunteers. As expected, anti-CHIPS IgG is detected in all 168 volunteers, resembling a Gaussian distribution [23] (Figure 3a). To limit undesired effects *in vivo*, only subjects with low anti-CHIPS titers were included in the study (antibody titer ≤ 3.92, as part of the exclusion criteria). To this end, we determined anti-CHIPS antibody titers in study subjects prior to receiving CHIPS. As expected, the 6 trail subjects have pre-existing anti-CHIPS antibodies (Figure 3a). Accordingly, anti-CHIPS IgG titers from subjects were within the normal range of tested sera, representative of the anti-CHIPS IgG titers of the general population (Figure 3a). The anti-CHIPS antibody titers in subjects were considered low enough to not affect the safety assessment of CHIPS.

**Fig 3:**
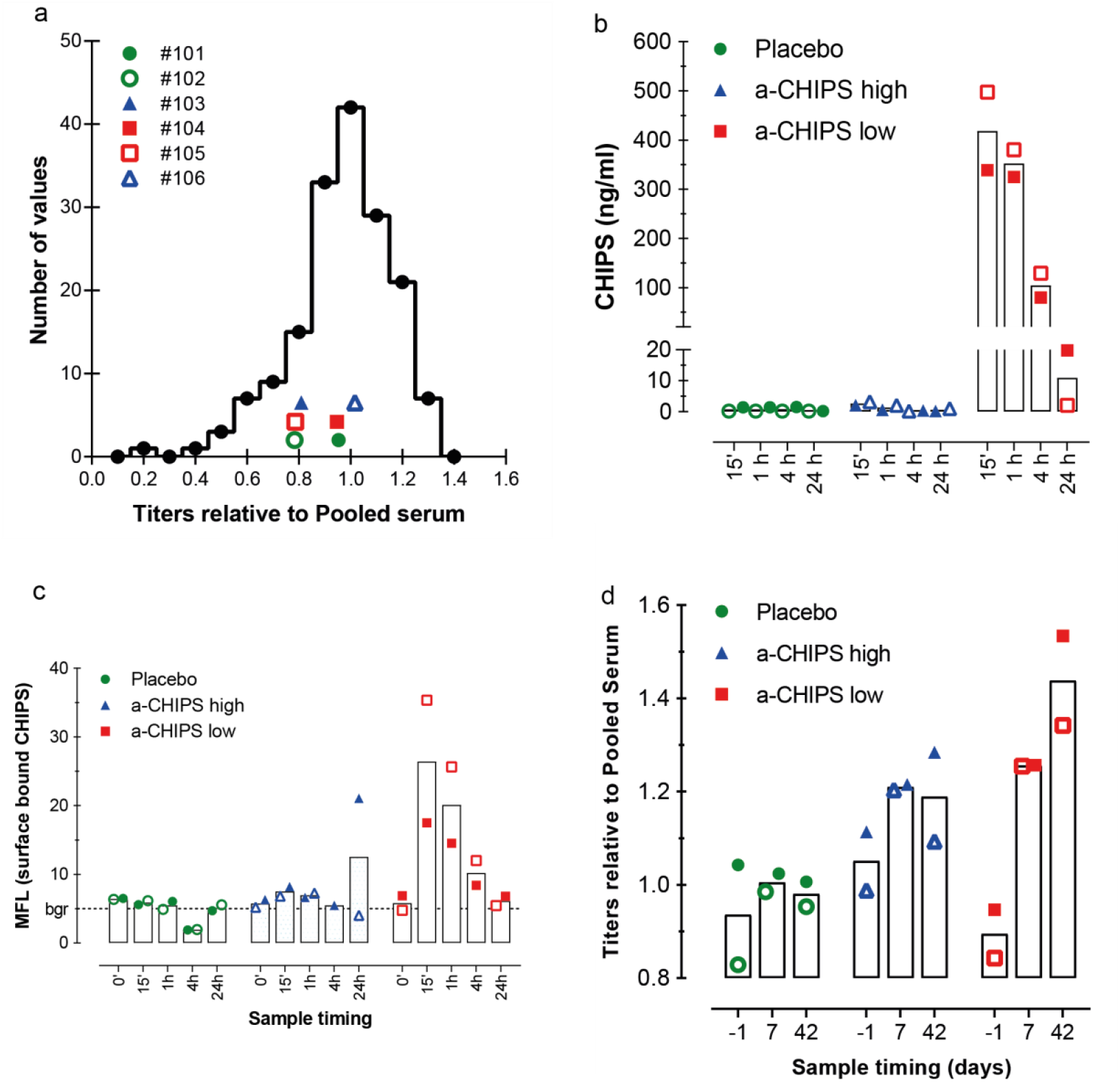
CHIPS and anti-CHIPS antibodies in humans. **a,** Frequency distribution of IgG anti-CHIPS titer in healthy human donors (n=168). The titer was defined as the log dilution that gives an absorbance of OD0.300 after subtraction of background value. Titers were depicted relative to the mean human pooled serum (HPS) titer (3.75). Anti-CHIPS antibody titer of the 6 subjects before study entry are depicted in the same graph as comparison. The ■ represents subjects that had low anti-CHIPS antibodies (anti-CHIPS low), ▲ represents subjects with high anti-CHIPS antibodies (anti-CHIPS high) and the ● represents subjects in the placebo group. Open and closed symbols differentiate between receivers in each group. **b,** Pharmacodynamics of CHIPS detected in the sera of the volunteers. CHIPS was measured by a specific capture ELISA at various time points after intravenous injection of CHIPS. **c,** CHIPS is recovered on the surface of peripheral blood neutrophils. At various time points after i.v. injection, the presence of CHIPS bound to the surface of neutrophils was detected with rabbit-anti-CHIPS antibodies. Values are expressed as mean fluorescence (MFL) of gated neutrophils in EDTA whole blood samples. Background MFL value for the secondary FITC labelled conjugated was 6. **d,** Immunogenicity of CHIPS in healthy human subjects. Specific IgG titer towards CHIPS were determined in all subjects before trial start, 7 and 42 days after trial closing and are depicted relative to HPS.

To further assess the safety of CHIPS, pre-clinical safety experiments were conducted in non-human subjects, prior to administration in humans. In all the animal toxicology studies, we did not observe any CHIPS-related toxicologically significant changes in clinical observations, body weight, food consumption, haematology, coagulation, blood chemistry parameters, ophthalmoscopy, electrocardiograms, macroscopic or microscopic pathology or behaviour (full pre-clinical assessment disclosed in supplementary text 1). Notably, a transient decrease in mean arterial blood pressure (40%) was observed in beagles receiving a high dose of 20 mg kg^−1^ CHIPS (supplementary text 1). However, mean arterial blood pressure returned to normal within 5 minutes post dosing. In all, these results suggest that side effects induced by CHIPS are unlikely to be observed in human subjects. As a result, the safety of CHIPS was subsequently studied in a set of six human subjects during a phase-1 clinical study.

Based on the toxicology studies, the administration of a single low dose of 0.1 mg kg^−1^ CHIPS was considered safe and administered in 4 human subjects. First, we determined the presence of CHIPS in sera of the volunteers during different time-points post-CHIPS administration. In only two out of four subjects that received the CHIPS protein, subjects #104 and #105, CHIPS could be detected 15 min post-i.v. injection with a gradual decline after 1 hour (Figure 3b). CHIPS was not detected in the sera of subjects #103 and #106 (Figure 3b). These observed differences in the detection of CHIPS in blood of the subjects seems to correlate with their initial level of anti-CHIPS antibodies. We hypothesized that the higher anti-CHIPS antibody titers hamper the detection of CHIPS by ELISA. Possibly, the epitope recognized by the capture monoclonal anti-CHIPS antibody is occupied by anti-CHIPS antibodies of the subjects. Consequently, we divided the 4 volunteers in 2 separate groups based on their anti-CHIPS antibody titer; anti-CHIPS low (subjects #104 and #105) and anti-CHIPS High (subjects #103 and #106). The measured CHIPS serum concentration in subjects #104 and #105 are also potentially an underestimation due to the interference of pre-existing anti-CHIPS antibodies. In addition, for subjects #104 and #105 that had detectable levels of CHIPS 15 min post i.v. injection, CHIPS concentrations dropped a 2-log fold over the course of 24 hours (Figure 3b). These data show that CHIPS is taken up systemic within 15 min and cleared after 24 hours post i.v. administration. We calculated a predicted half-life of CHIPS to be at least 1.5 hours in humans.

CHIPS binds the C5aR1 on human neutrophils with high affinity *ex vivo* [3]. However*, in vivo* binding of CHIPS could be hampered by circulating antibodies. In order to assess if CHIPS interacts with its cellular target, we determined the binding of CHIPS *in vivo* on neutrophils of the subjects. The presence of CHIPS on the surface of neutrophils was determined at various time points post-CHIPS administration using a rabbit-anti-CHIPS antibody [27]. Notably, the binding of CHIPS on the surface of neutrophils was only detected in subjects with a low anti-CHIPS antibody titer (subjects #104 and #105) (Figure 3c). It is possible that the circulating anti-CHIPS antibodies present in serum also interfere with the direct detection by the specific anti-CHIPS monoclonal antibody or even the direct association with the C5aR on neutrophils. Therefore, the lack of a direct detection cannot exclude the absence nor presence of CHIPS bound to the receptor in the individuals with high anti-CHIPS antibody titers. All in all, we show that CHIPS binds circulating human blood neutrophils, confirming the interaction with target cells *in vivo*.

All tested subject had pre-existing anti-CHIPS antibodies. As a specific antibody response is mediated against CHIPS, it is likely that a re-challenge with CHIPS will lead to an increase in antibody titers. To determine the immunogenicity of CHIPS, anti-CHIPS serum titers were measured during different time points pre- and post-CHIPS administration. An increase in anti-CHIPS titer was observed in individuals receiving CHIPS that had a low anti-CHIPS antibody titer (subjects #104 and #105) pre-CHIPS administration (Figure 3d).The rapid boost of circulating IgG titers by the staphylococcal protein CHIPS in humans indicates high immunogenicity and pre-existing memory, supporting a concept of expected exposure to secreted staphylococcal proteins starting at early age [24,25,28].

### CHIPS induced adverse effects in humans

The administration of CHIPS in human subjects was tolerated by 2 subjects (subjects #103 and #104), moderately tolerated in subject #105 but subject #106 (subject with a high anti-CHIPS antibody titer, open blue triangle in figures) developed serious symptoms directly after the CHIPS infusion, which were diagnosed as an anaphylactic reaction (Supplementary text 2). No adverse events were reported in subjects receiving placebo. To determine whether the subjects developed a CHIPS-mediated inflammation response, white blood cell count (WBC) and C-reactive protein concentration (CRP) were measured pre- and post-dosing. CHIPS induced a transient leukocytopenia in the subjects receiving CHIPS that resolved within 2 days (Figure 4a). Within the group of subjects that received CHIPS there was a mild increase in CRP (average of 42 mg ml^−1^) at day 2 post CHIPS dose compared to controls. CRP levels returned to normal when subjects were screened during follow up at day 15 (Figure 4b). This indicates that there was indeed an inflammation response upon CHIPS administration.

**Fig. 4:**
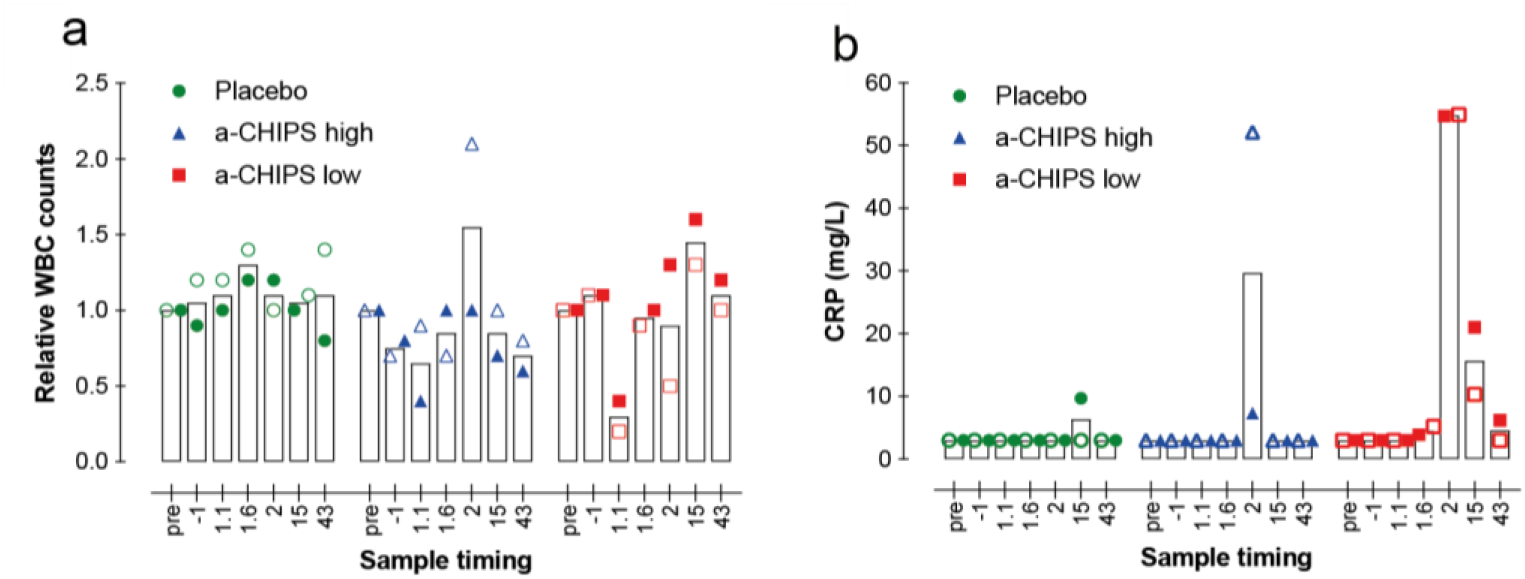
CHIPS induces leukocytopenia and increased CRP levels in humans. Levels of circulating **a,** peripheral white blood cells and **b,** serum inflammation marker CRP. At various time points after intravenous injection of CHIPS, WBC counts and CRP measurements were performed. (1.1 and 1.6 indicate 1 day and 1 or 6 hours respectively). Data for WBC are expressed relative to the value at T = 0 and data for CRP are expressed in mg/mL. The ■ represents subjects that had low anti-CHIPS antibodies (anti-CHIPS low), ▲ represents subjects with high anti-CHIPS antibodies (anti-CHIPS high) and the ● represents subjects in the placebo group. Open and closed symbols differentiate between receivers in each group.

### Circulating immune complexes and increased serum tryptase

Mast cells play a central role in anaphylaxis and other allergic conditions. Immune complexes can activate mast cells by FcR crosslinking and through activation of complement and the generation of C5a [29]. Circulating immune complexes (CIC) induce the abundant secretion of the serine proteinase tryptase by mast cells, which can be used as an indicator of anaphylaxis. Since all subjects had pre-existing anti-CHIPS antibodies, we evaluated whether intravenous administration of CHIPS leads to the formation of CIC. Circulating immune complexes were detected in the subjects receiving intravenous CHIPS (Figure 5a). Subject #106, who suffered an anaphylactic reaction following the administration of CHIPS, showed the highest CIC levels, contrary to subjects #104 and #105 who remained at baseline. CIC were also detected in subject #103, who has the highest anti-CHIPS antibody titer but reported only minor adverse effects. No CIC were detected in subjects that received the placebo.

**Figure 5:**
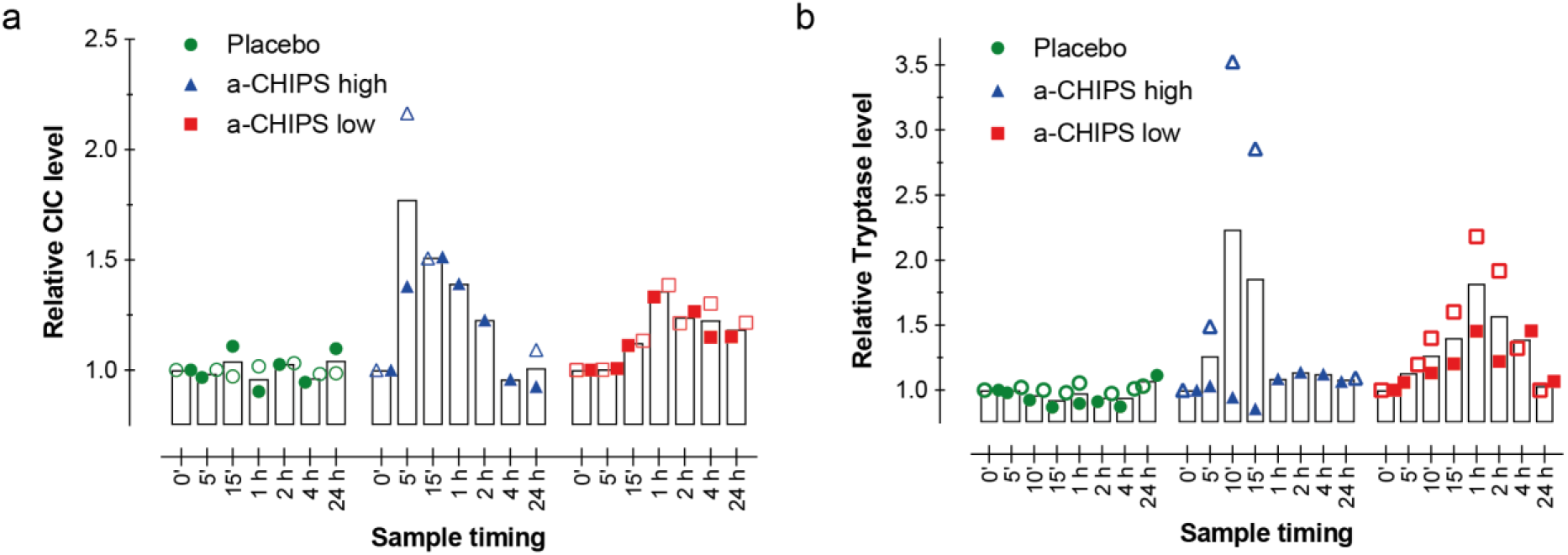
Circulating immune complexes and increase serum tryptase. Adverse effects of CHIPS as measured by levels of **A)** Circulating Immune Complexes (CIC), and **B)** mast cell marker tryptase. At various time points after intravenous injection of CHIPS, specific assays were performed for both markers. The ■ represents subjects that had low anti-CHIPS antibodies (anti-CHIPS low), ▲ represents subjects with high anti-CHIPS antibodies (anti-CHIPS high) and the ● represents subjects in the placebo group. Open and closed symbols differentiate between receivers in each group.

Subsequently, we measured the serum tryptase levels in the subjects. An increase in serum tryptase concentration was detected in all subjects receiving CHIPS except subject #103, that reached a maximum at approximately 10 minutes post dose and continued to drop to baseline levels after 24h (Figure 5b). Notably, subject #106 had the highest levels of tryptase, which correlates with the high levels of CIC measured. These data suggest that CHIPS administration in subjects with high circulating anti-CHIPS titers results in an inflammatory response and adverse effects. Due to these effects, the study was stopped and no further administrations of CHIPS was undertaken.

## Discussion

In this study we show that the staphylococcal secreted CHIPS as a model virulence factors-based therapeutic agent, is not suitable for systemic use in humans. This is due to the presence of pre-existing circulating CHIPS antibodies present in humans. Despite the neutralizing effect of anti-CHIPS antibodies, we were able to detect significant serum concentrations of the CHIPS protein. The development of a second-generation CHIPS protein with a preserved activity but with reduced immunogenic properties could make a promising new candidate anti-inflammatory drug. Mapping of the epitopes for human IgG within the CHIPS protein will be an important first step in this development [30]. We previously identified several unique conformational epitopes on CHIPS using affinity purified human IgG from a preparation for intravenous use [30]. However, despite developing a version of CHIPS with low interaction with pre-existing anti-CHIPS antibodies, the high immunogenicity of CHIPS will limit it suitable for therapies requiring a single administration. All in all, the development of a human C5aRKI mouse made it possible to asses CHIPS as a therapeutic agent in C5aR mediated diseases. However, human C5aRKI mice could also be used to assess CHIPS as a virulence factor and better understand the contribution of CHIPS to Staphylococcal pathophysiology. The use of our human C5aRKI mouse has already contribute to our understanding of staphylococcal pathophysiology by elucidating the in vivo role of the human C5aR interacting staphylococcal bi-component toxin HlgCB [19].

Besides CHIPS, other staphylococcal proteins have been suggested as potential therapeutic agents for a variety of inflammatory diseases. Previous studies with staphylococcal proteins that intervene with C5 complement activation (SSL7 and Ecb) or the FcγR (FLIPr-Like) also proved to be effective inhibitors in a murine Arthus model [21,31,32]. However, as pre-existing circulating antibodies against many if not all staphylococcal immune evasion proteins are present in all human [26], the use of these staphylococcal proteins as therapeutic agents are most likely severely hampered. In addition, antibodies and neutralizing activity against staphylococcal virulence factors can last up to six months post-administration [33]. Therefore, even if the primary dose is tolerated, re-administration should be avoided. Despite the drawbacks of using staphylococcal immune evasion molecules, other bacterial virulence factors have shown to be possibly applicable as therapeutics. The *S. pyogenes* virulence factor Immunoglobulin G-degrading Enzyme of *S. pyogenes* (IdeS) ablates the humoral immunity by cleaving and inactivating IgG [34]. Even though humans carry anti-IdeS antibodies, IdeS treatment also effectively neutralizes IdeS-specific IgG [35]. IseD was suggested as a way of helping at preventing antibody-mediated injury to allografts. During the combined phase 1 and 2 trials, however, a total of 38 serious adverse effects in 15 patients were witnessed [36]. The use of IdeS did consistently reduced or eliminated donor-specific antibodies to desirable levels, allowing transplantation form an HLA-incompatible donor [36]. Although bacterial immune evasion molecules are not suited for direct use as therapeutic compounds, future molecules based on the bacterial anti-inflammatory proteins could very well be potential new candidates. Knowledge of the exact mechanisms of action and the active sites can lead to the development of small molecule anti-inflammatory drugs based on bacterial virulence factors.

## Materials and Methods

### Ethics statement

The RCT study protocol (JPD-003/002/NL) and amendments were approved by an independent ethics committee. The study was performed in compliance with the ‘Declaration of Helsinki’ (Scotland, October 2000) and OECD Principles of Good Laboratory Practice and applicable regulatory regulations. For neutrophil isolation approval was obtained from the medical ethics committee of the University Medical Center Utrecht (METC‐protocol 07‐125/C approved March 01, 2010; Utrecht, The Netherlands). The use of animals was approved by the National Ethical Committee for Animal Experiments and performed according to the guidelines of the Central Animal Facility of the Utrecht University (Project# AVD115002016565).

### Isolation of Rabbit anti-ovalbumin IgG

IgG was purified from Rabbit anti-Chicken-Egg Albumin, delipidized whole antiserum (Sigma-Aldrich) using multiple runs over a 1 ml Protein-A HiTrap column (GE Healthcare Life Sciences) on an ÄKTA FPLC (GE Healthcare Life Sciences). Rabbit IgG was eluted from the column with 0.1 M citric acid, pH 3.0 and collected fractions were neutralized with 1M Tris-HCl, pooled and dialyzed against PBS. Protein concentration was determined at 280 nm using a molar extinction coefficient of 1.35 for Rabbit IgG.

### Peritoneal Arthus reaction and neutrophil migration

Human C5aR1^KI^ mice were generated and characterized as described elsewhere [19]. The Arthus reaction was initiated upon i.v. injection in hC5aR1^KI^ mice (male and female) of 100 μl of OVA (20 mg kg^−1^ of body weight; Sigma-Aldrich) immediately followed by an i.p. injection of 800 μg of rabbit anti-OVA IgG (Sigma) in 500 μl PBS. For mice in the CHIPS group, 60 μg CHIPS was administered i.p. 30 minutes prior to initiation of the Arthus reaction and simultaneously with OVA i.v.. For the control group, PBS was administered i.v. and i.p.. Mice were euthanized by CO_2_ suffocation 6-hours after the onset of the peritoneal Arthus reaction and the peritoneal cavity washed with two times 5 ml of ice-cold RPMI 0.1% HSA/5mM EDTA. Peritoneal fluid was recovered and centrifuged at 1200 rpm for 10 min to collect the exudate cells. Cell pellets were resuspended in 500 μl buffer and counted with trypan blue in a TC20 automated cell counter (BioRad). Cells were stained in the presence of a Fcγ-receptor blocker, with anti-mouse CD45-APC (clone 30-F11, BD Biosciences), anti-mouse Gr1-PE (1A8, BD Biosciences), anti-mouse F4/80 FITC (BM8, eBioscience), anti-human C5aR-FITC (clone S5/1, SeroTec), isotype rat-IgG2a-FITC (R&D) and rat-IgG2b-PE (BD Biosciences). Samples were analyzed by flow cytometry. Collected peritoneal cells were washed with PBS and the cell number adjusted to 5×10^6^ cell ml^−1^. Cytospin slides were prepared with 50 μl 5×10^4^ cell suspension and stained with DiffQuick. The percentage of neutrophils was determined by flow cytometry analysis and confirmed by the number of neutrophils based on morphology following DiffQuick staining. Mouse neutrophils were isolated from bone-marrow as described elsewhere [37,38]. Briefly, bone marrow cells were collected by flushing the femurs and tibias with 10 ml of cold HBSS + 15 mM EDTA + 30 mM Hepes + 0.1 % HSA. A two-layer Percoll density gradient (2 ml each in PBS) composed of 81% and 62.5% was used to enrich neutrophils from the total leucocyte population. Interphase between 62.5% and 81% was collected. Cells were washed once with buffer and resuspended in PRMI1640 with 0.1% HSA. Staining of bone marrow cells was performed as described above.

### Preclinical assessment of CHIPS toxicity in animal models

Conventional pre-clinical toxicology studies were preformed to investigate the safety of intravenous CHIPS. These included; (I)The effects of CHIPS on various cardiovascular and respiratory parameters in one group of three anesthetized beagle dogs. The dogs were administered CHIPS in incremental doses of 0.2, 2.0 and 20 mg kg^−1^, infused intravenously over 1 minute at approximately 30 minute intervals. (II) Behavioral (’Irwin’) test in mice: CHIPS was administered as a single intravenous injection to male ICR CD-1 mice (3 per group) at doses of 7.5, 25 and 75 mg kg^−1^ in order to assess effects on general behavior. An additional group received an equivalent volume (10 ml kg^−1^) of vehicle (0.9% w/v sterile saline). (III) Acute intravenous toxicity study in rats: Intravenous administration of 96.1 mg∙kg^−1^ CHIPS as a single dose (the maximum practically achievable due to volume considerations) to 5 male and 5 female rats. (IV) Acute intravenous toxicity in mice: Intravenous administration of 96.1 mg kg^−1^ CHIPS as a single dose to 5 male and 5 female mice. (V) Seven day intravenous bolus preliminary toxicity study in rats (24 males and 24 females, maximum dose 10 mg kg^−1^). (VI) Seven day intravenous bolus toxicity study in rats (76 males and 76 females, maximum dose 10 mg kg^−1^). (VII) Seven day intravenous bolus dose range finding study in dogs (2 males and 2 females, maximum dose 20 mg kg^−1^). (VIII) Seven day intravenous bolus toxicity study in the dogs (12 males and 12 females, maximum dose 20 mg kg^−1^).

### Inclusion of human volunteers

Full description of study population, including number of subjects, inclusion, exclusion and removal criteria are described in supplementary Protocol No.: JPD-003/002/NL. Briefly, inclusion criteria for healthy volunteers were as follows: (I) Adult males within an (II) age range 18-50 and (III) a body mass index (BMI) of 18-30 kg m^−2^. Medical screening was divided in 2 parts. Subjects were screened for anti-CHIPS antibody titers. Only subjects with a low titer (equal or lower to 3.92, defined as the log of the serum dilution that gives an absorbance value of 0.300 in the ELISA) were screened for the second part within 3 weeks before dosing and include: medical history, physical examination, measurement of blood pressure, heart rate, respiration and temperature, alcohol breath test, blood and urine tests, electrocardiogram (ECG) and drug screening.

### Admission and follow up

Full description of the admission and follow up, treatments and stopping rules are described in (supplementary Protocol No.: JPD-003/002/NL). Briefly, six selected subjects (4 receiving CHIPS and 2 controls) were admitted to the Clinical Pharmacology Unit (Kendle, Utrecht, The Netherlands) on the day before dosing. Baseline measurements, including blood samples for safety, urinalysis, interim medical history, physical examination, vital signs and ECG were done. On the day of dosing CHIPS (0.1 mg kg^−1^ administered as a single dose of sterile frozen isotonic saline solution containing CHIPS at a concentration of 5 mg ml^−1^) or placebo (0.9% NaCl) was administered by intravenous infusions over 5 minutes. Subjects were connected to a telemetry system for cardiac monitoring from 30 minutes before dosing until 4 hours after start of dosing. The blood pressure of subjects was measured continuously using a Finapres from 5 minutes before dosing until 30 minutes after start dosing. Vital signs were measured and ECG’s were made at certain time points during the admission period. For safety, clinical status and laboratory values (haematology, biochemistry, coagulation and urinalysis) of all subjects were monitored. Adverse events were documented and characterized according to their severity and relationship to CHIPS or placebo. The subjects were discharged at 24 hours after dosing. Two weeks after dosing subjects returned to the Unit for a visit to evaluate vital signs, ECG, blood and urine and anti-CHIPS antibody level. A follow up visit was scheduled 6 weeks after dosing.

### Cloning and expression of CHIPS

CHIPS was cloned and expressed as described earlier [3,27]. Briefly, the CHIPS gene (*chp*; GenBank: AF285146.1), without the signal sequence, was cloned into the pRSET vector directly downstream the enterokinase cleavage site and before the EcoRI restriction site by overlap extension PCR. Bacteria were lysed with CelLytic B Bacterial Cell Lysis/Extraction Reagent (Sigma) and lysozyme according to the manufacturer’s description. The histidine-tagged protein was purified using a nickel column (HiTrap Chelating HP, 5ml, Amersham Biosciences) following the manufacturer’s instructions and cleaved afterwards with enterokinase (Invitrogen). Samples were checked for purity and presence of protein using 15% SDS-PAGE (Polyacrylamide gel electrophoresis, Mini Protean 3 System, Bio-Rad) and Coomassie Brilliant Blue (Merck) staining.

### Purification of CHIPS for intravenous use

Full-length CHIPS was expressed in *E. coli* containing the coding sequence of CHIPS directly downstream to PelB coding sequence in a growth media consisting of soya peptone and yeast extract in 8 liter fermentation media. CHIPS was isolated both from the growth media and the cells by a two stage cation exchange purification process followed by a desalting step. The bacterial cell pellet was resuspended in phosphate buffer (30 mM; pH 7.0), containing NaCl (10 mM), DTT (10 mM) and frozen. This was subsequently thawed at 37°C, incubated on ice and sonicated. After centrifugation at 15,000 rpm, an amber colored ‘cell’ supernatant was recovered. The supernatant was diluted four-fold with 30 mM phosphate buffer and passed over a Source S-30 column. The material was eluted with a phosphate buffer salt gradient and fractions containing CHIPS were combined and purified further by using a polishing column with a shallow salt gradient. Fractions containing CHIPS with purity greater than 97% (by HPLC) were combined and passed through a Sephadex G 25 desalting column to remove phosphate and excess of sodium chloride. Endotoxin was removed by gently shaking over resin (Biorad) and the preparation was sterilized through ultra-filtration. We confirmed the purity by HPLC-MS on a Microbondapac CN-RP column with a mobile gradient phase consisting of water-TFA to Methanol-TFA. The end product was diluted with sterile saline to the desired concentration and stored at −20°C.

### Isolation of human PMN

Blood obtained from healthy volunteers was collected into tubes containing sodium heparin (Greiner Bio-One) as anticoagulant. Heparinized blood was diluted 1/1 (v/v) with PBS and layered onto a gradient of 10 ml Ficoll (Amersham Biosciences, Uppsala, Sweden) and 12 ml Histopaque (density 1.119 g ml^−1^; Sigma-Aldrich, St. Louis, MO). After centrifugation (320 g, for 20 min at 22°C), the neutrophils were collected from the Histopaque phase and washed with cold RPMI 1640 medium containing 25 mM HEPES buffer, L-glutamine (Invitrogen Life Technologies) and 0.05% HSA (Sanquin, Amsterdam, the Netherlands). The remaining erythrocytes were lysed for 30 s with ice-cold water, after which concentrated PBS (10 x PBS) was added to restore isotonicity. After washing, cells were counted and resuspended in RPMI-1640 / 0.05% HSA at 10^7^ neutrophils ml^−1^.

### Determining Circulating Immune Complexes, C-Reactive Protein and serum tryptase

CIC were determined by 2 different ELISA’s from Quidel (San Diego, CA): the CIC-C1q enzyme immunoassay is based on the principle that complement fixing IC will bind to immobilized human C1q purified protein; the CIC-Raji Cell Replacement enzyme immunoassay measures IC containing C3 activation fragments by using a mAb that specifically binds the iC3b, C3dg and C3d activation fragments of C3 in a manner which is analogous to the classical Raji cell CR2 binding reaction. The data of both assays were combined and results expressed relative to the value at time point 0. CRP levels were determined by the diagnostic department according to standard protocols. Serum derived tryptase (both α-and β-form) was measured on the UniCAP®-100 using the ImmunoCAPTM technology (Pharmacia Diagnostics, Woerden, The Netherlands). The normal geometric mean for serum tryptase in healthy controls is 5.6 μg l^−1^. Results were expressed relative to the value at time point 0.

### ELISA for anti-CHIPS antibodies and CHIPS levels

Rabbits were immunized with recombinant CHIPS using Freund’s Complete Adjuvants and boosted with Freund’s incomplete adjuvants. Bleedings were checked for reactivity with CHIPS by ELISA as described for human anti-CHIPS antibodies (see below). From the final bleeding, IgG was purified by standard Protein-G (Pharmacia) affinity chromatography according to the manufacturer’s instructions. For the anti-CHIPS ELISA, microtiter plates (Greiner) were coated with 50 μL CHIPS per well at 1 μg ml^−1^ in PBS overnight at 4°C. All wash steps were performed thrice with PBS-0.05%Tween-20 and subsequent incubations were done for 1 hour at 37°C. Plates were blocked with PBS-0.05%Tween-20 4% BSA, washed and incubated with sera or antibodies diluted in PBS-0.05%Tween-20 1% BSA. Bound antibodies were detected with species-specific goat anti-IgG conjugated with peroxidase (all from Southern, Birmingham, USA) and TMB as substrate. The reaction was stopped with H_2_SO_4_ and the absorbance measured at 450nm in a BioRad ELISA-reader. For the capture ELISA, microtiter plates were coated with 50 μL α-CHIPS mAb 2G8 at 3 μg mL^−1^ in PBS overnight at 4°C. Plates were blocked with PBS-0.05%Tween-20 4% BSA, washed and incubated with diluted samples and a two-fold dilution range of CHIPS as standard in PBS-0.05%Tween-20 4% BSA. Subsequently, plates were incubated with 0.33 μg mL^−1^ rabbit α-CHIPS IgG and 1:5000 diluted peroxidase-conjugated goat anti-rabbit IgG (Southern). Bound antibodies were quantified with TMB as substrate, the reaction stopped with 1 N H_2_SO_4_ and OD was measured at 450 nm on a BioRad ELISA reader.

### Statistical analysis

Calculations of statistical analyses were performed using Prism 7.0 (GraphPad Software). Flow cytometric analyses were performed with FlowJo (Tree Star Software). Significance was calculated using analysis of variance (ANOVA) followed by Kruskal-Wallis as post-test correction for multiple comparison. All statistical methods with regards to the human trails are described in the supplementary (Protocol No.: JPD-003/002/NL.)

## Acknowledgements

We thank Miriam J.J.G. Poppelier, Miranda Boonstra, Toon van Bommel and Maroeska Oudshoorn (University Medical Center Utrecht, Utrecht, the Netherlands) for their technical support.

## Supplementary Text 1: Pre-clinical assessment of CHIPS as a therapeutic agent

CHIPS successfully dampened the C5aR dependent Arthus reaction in a mouse model expressing the hC5aR. To allow testing of CHIPS in human volunteers, pre-clinical safety experiments in non0hman subjects were required. In none of the toxicology animal studies did administration of CHIPS cause any CHIPS related toxicologically significant changes in clinical observations, body weight, food consumption, haematology, coagulation, blood chemistry parameters, ophthalmoscopy, electrocardiograms, macroscopic or microscopic pathology or behavior (Data?). The effects of CHIPS on various cardiovascular and respiratory parameters in anesthetized beagle dogs was examined. In the dogs receiving low dose CHIPS (0.02 and 2 mg‧kg-1) there was no evidence of cardiovascular or respiratory effects when compared to infusion of vehicle (isotonic saline) (Data?). Following intravenous administration of 20 mg‧kg-1 CHIPS a transient decrease in mean arterial blood pressure (40%) was recorded approximately 1 minute after start of administration(Data?). Mean arterial blood pressure levels returned to pre-dose levels within approximately 5 minutes following the start of dosing (Data?). The effect on blood pressure coincided with transient, inconsistent changes in heart rate. One dog was administered a repeat intravenous dose of CHIPS (20 mg‧kg-1) approximately 30 minutes following the first administration of CHIPS. Transient effects on cardiorespiratory parameters similar to those recorded following the first dose were not apparent after the repeat administration of CHIPS. However, the second administration produced a prolonged reduction in mean arterial blood pressure, reaching a maximum of 18% at approximately 30 minutes following the second administration(Data?). In this animal only, twelve minutes following the repeated administration of CHIPS a generalized skin reaction appeared consistent with some form of mild allergic reaction. The results of this study suggested that cardiorespiratory effects are unlikely to be observed in the human subjects in the used dose range (0.1 mg‧kg-1). Furthermore, any effects that might occur were expected to be transient and reversible.

